# Quantifying the IUCN Red List: Using historical assessments to calculate future extinction risk

**DOI:** 10.1101/2024.02.04.578808

**Authors:** Ryan Bates, Elizabeth Taylor, Yuheng Sun, Rikki Gumbs, Monika Böhm, Claudia Gray, James Rosindell

## Abstract

The IUCN Red List of Threatened Species assigns species to discrete categories based on a common set of criteria to capture relative extinction risk. Some quantitative analyses require a continuous probability of extinction not provided by discrete categories. Furthermore, the criteria may not translate to extinction risk in a universal way across species, because some species’ attributes are not incorporated into the criteria. Here we calculate extinction risk using Markov Models parameterised with historical IUCN Red List assessment data, and we provide quantitative extinction risk for use in downstream analyses. We find that taxonomy, body size, and habitat specialism interact with extinction risk in ways not fully captured by the existing categories. For example, an Endangered habitat specialist appears equally at risk compared to a Critically Endangered habitat generalist. We hope our findings will support more accurate quantitative biodiversity analyses of species extinctions and declines using Red List data.

## 1 Introduction

We are in the middle of a biodiversity crisis on par with past mass extinction events (Barnosky et al. 2011; Ceballos et al. 2015; McCallum 2015). Many organisations - such as the IUCN (International Union for the Conservation of Nature) and WWF (World Wide Fund for Nature) - are measuring declines in biodiversity across the globe. Their work details the nature of biodiversity declines and advises on ways to slow or reverse them. For example, the Living Planet Index and Red List Index (WWF 2022; S. H. Butchart et al. 2004) are both used as progress indicators for the Convention on Biological Diversity’s new Kunming-Montreal Global Biodiversity Framework (GBF) (Convention on Biological Diversity 2022). Many aspects of the GBF relate to extinction risk. For example, the GBF Goal A asks for a tenfold reduction of extinction rate and risk by 2050 (CBD 2022). To check if we are on target to meet this goal, as well as many others, requires global-scale quantitative measures of extinction risk.

A global indicator for extinction risk used in many studies is the IUCN Red List of Threatened Species (hereafter Red List). There are six Red List categories for extant species with sufficient data: Least Concern (LC), Near Threatened (NT), Vulnerable (VU), Endangered (EN), Critically Endangered (CR), and Extinct in the Wild (EW). These categories were initially proposed by Mace and Lande (1991) and fully implemented in 2001 (IUCN 2001). Species in VU, EN and CR categories are considered ‘threatened’ with extinction. Species are assessed for the Red List using five criteria (A-E), and are placed in the highest risk category triggered by any criteria (IUCN 2012). These criteria cover various key indicators of a species’ extinction risk, from population size to spatial distribution.

Criterion E is the only criteria directly based on probability of extinction. It enables quantitative extinction risk values, often coming from Population Viability Analyses, to be mapped to the three threatened categories (IUCN 2012).

The Red List has not only been applied as the cornerstone of indicators, such as the Red List Index, it has also been used extensively in studies seeking to investigate which ecological or behavioural attributes increase species’ risk of extinction (for example see Cardillo et al. 2004; Ducatez and Shine 2017; Kotiaho et al. 2005; Purvis, Agapow, et al. 2000; Purvis, Gittleman, et al. 2000; Webb and Mindel 2015). Such studies typically compare threatened to non-threatened species, or use the discrete Red List categories as the starting point of quantitative analyses. Using all six categories of the Red List as opposed to comparing threatened and non-threatened species allows for more detailed analyses and findings. Nevertheless, simply numbering categories limits the quantitative gap between them to a linear (or log-linear) relationship between the extinction risk and Red List category. Such a restriction does not appear consistent with the quantitative gaps between Red List category criteria. For example, under criterion B2 the maximum area of occupancy to qualify for certain Red List categories is 2000km^2^ for VU, 500km^2^ for EN, and 10km^2^ for CR (IUCN 2012), which is neither linear nor log-linear. A more nuanced relationship between Red List category and risk of extinction over a given timescale seems likely, yet few studies have attempted to quantify this. A further complication of using the Red List to investigate how species’ traits correlate with extinction risk is that any such analysis assumes the Red List categories are universal; i.e. a random Endangered species should be no more or less at risk than any other random Endangered species, regardless of taxonomy or other species’ attributes. The Red List criteria implicitly account for some effects, for example body mass via generation length and population size, which are known to correlate with body mass (Currie 1993). Yet it remains a challenge to investigate which further species attributes might be taken into account to enhance the accuracy of our extinction risk estimates. If ecological factors affect extinction risk in ways that are not fully captured by Red List category, then this may lead to a bias in downstream analyses.

A method to convert Red List categories into quantitative extinction risk and check their consistency across species would have numerous applications. Firstly, investigations of the drivers of extinction risk would represent the true risk more accurately, and on a continuous scale. Secondly, key indicators could be interpreted as an approximation of extinction risk over a given timescale, rather than relative extinction risk or a proxy as is the case currently. Finally, a suite of conservation prioritisation schemes such as EDGE2 and HEDGE (Isaac et al. 2007; Steel et al. 2007) already require probabilities of extinction explicitly by their construction.

Criterion E, which is directly based on probability of extinction, has previously been extrapolated in order to estimate extinction probabilities for all five extant Red List categories over various time-frames (Mooers et al. 2008). Specifically, the approach maps the Red List categories to an exponential pattern of extinction risk, assuming that all other criteria (A through D) translate to the same probability of extinction as criterion E (Mooers et al. 2008). This assumption was necessary to get a probability of extinction with a quantitative rationale. However, it is questionable because the IUCN deliberately chose to specify criterion E separately rather than to frame it as a descriptor of extinction risk that would apply to species assessed under criteria A-D (Mace, Collar, et al. 2008).

An explicit model of historical changes in species’ Red List categories over time provides a potential solution that would avoid relying on criterion E. Previous studies have applied a multi-state Markov (MSM) model to predict movement between Red List categories, including into the Extinct category, but were restricted to using bird data (Andermann et al. 2021; Monroe et al. 2019). Monroe *et al*. (2019) found that extinction risks were an order of magnitude or more below the estimates based on Criterion E, a finding with potentially large ramifications for downstream calculations using probabilities of extinction. Monroe’s findings were based on Red List Index data for birds (Monroe et al. 2019). Red List Index data are a small subset of the full Red List that are particularly suitable for such analyses as they provide historical Red List assessments which are amended or backcast, currently consisting of birds, mammals, amphibians, cycads and corals (S. H. M. Butchart et al. 2007). Backcasting is necessary to account for non-genuine changes in Red List category, e.g. changes caused by taxonomic revision or previous lack of data. Since only genuine changes are retained in the Red List Index data, they provide the cleanest view of movements between categories for the focal species. A study of extinction risk using historical movements between Red List categories across a broader range of taxa would be a valuable step. However, a major challenge in doing this is that Red List Index data is very limited in taxonomic scope whilst Red List data has not been backcast.

Here we aim to quantify probability of extinction both across and within Red List categories, using the full set of historical Red List assessments. We develop a new cleaning procedure using validity data from elsewhere in the IUCN Red List to filter transitions between Red List categories. This is not backcasting, but it still mitigates to some extent the effect of non-genuine changes. We analyse these clean data with MSM models to predict future changes in IUCN threat category (including to extinction), built using historical IUCN assessment data. The MSM model predicts extinction probabilities at any given point in the future, given a species’ current Red List assessment category. Our technique not only allows the extinction probabilities of Mooers *et al*. (2008) to be checked against real-world data, but also enables us to investigate the impact of other attributes on the probability of extinction within any given Red List category. We also investigate the probabilities of extinction when placing CR(possibly EX) and CR(possibly EW) species in the Extinct category. Given data availability and ecological factors already known to impact extinction risk (Chichorro et al. 2022; Purvis, Gittleman, et al. 2000), we consider the effects of higher taxa, body size and habitat specialisation on extinction risk. Does an Endangered bird species with generalist habitat preferences have a lower extinction risk than a bird species with specialised habitat requirements that is also Endangered? Does a Vulnerable plant have the same extinction risk as a Vulnerable mammal? Through our analysis we answer these questions and investigate how species attributes may have an impact on extinction risk even when comparing species within the same Red List category. Generating quantitative extinction probabilities is the key to understanding the patterns of extinction risk across all species, and understanding which species attributes may impact this risk. We hope our updated method of assigning probabilities of extinction to Red List categories will facilitate new quantitative analyses and provide more robust biodiversity indicators.

## 2 Materials and Methods

We acquired data from the Red List API using the R package rredlist (Gearty and Chamberlain 2022). These data included both species data and historical Red List assessment data from 1994 onward (correct as of Dec 2021). We downloaded the ‘Table 7 data’ from the IUCN Red List’s website which tells if past changes in any species’ Red List category were ‘genuine’ (due to real world changes in the species) or ‘non genuine’ (due to changes in information availability). We converted Table 7 (hereafter validity data) from PDF to a usable table format using Adobe’s free software (Adobe 2021).

All data processing was carried out in R (R Core Team 2023). We first ensured that only assessments made using the current criteria (V3.1 - IUCN 2012) remained in the data. Next, we merged EW assessments with EX assessments as very few assessed species were assigned EW (111 out of 114880). EX was chosen over CR due to the low rates of successful reintroduction (Smith et al. 2023), and the fact that EW species are no longer impacting native ecosystems. The Red List data lists CR(PEW) and CR(PEX) species as CR, so we were unable to look at the potential implications of these categorisations for the data as a whole. Merging EW species into EX was necessary to ensure there were enough examples of transitions into and out of each remaining category to enable the model to be parameterised. We incorporated validity data (see supplementary materials) to correct for non-genuine transitions; where previous assessments were based on incomplete evidence, or invalidated by taxonomic reassignment. The validity data also allowed us to handle Data Deficient (DD) species as DD species do not inherently have a threat status (See supplementary materials 4). Following this, we removed species with only one genuine assessment, as the modelling steps require a minimum of two assessments per species. Mindful that the validity data we use (from Table 7) was not intended by IUCN to be used in this way, we also repeated our analyses for Birds using Red List Index data, and using historical Red List data with no attempt to correct for validity for comparison to our main results.

We built a Multi-State Markov model (MSM model) using the package msm (Jackson 2011) in R. The model assumes that species must travel sequentially between states; e.g. a species cannot jump from LC to EN, and if that transition is seen in the data it is assumed they must have travelled through NT and VU without those being observed (as shown in Figure 1). The model also assumes that species cannot recover from becoming extinct, treating it as an absorbing state (Figure 1, more info in Supplementary Information). We modelled the historical Red List assessment data with the MSM model 100 times, each time randomly sampling the full data with replacement (bootstrapping) to generate uncertainty bounds. We capped the model output at 100 years, the maximum timescale used by Red List Criterion E (IUCN 2012).

**Figure 1:**
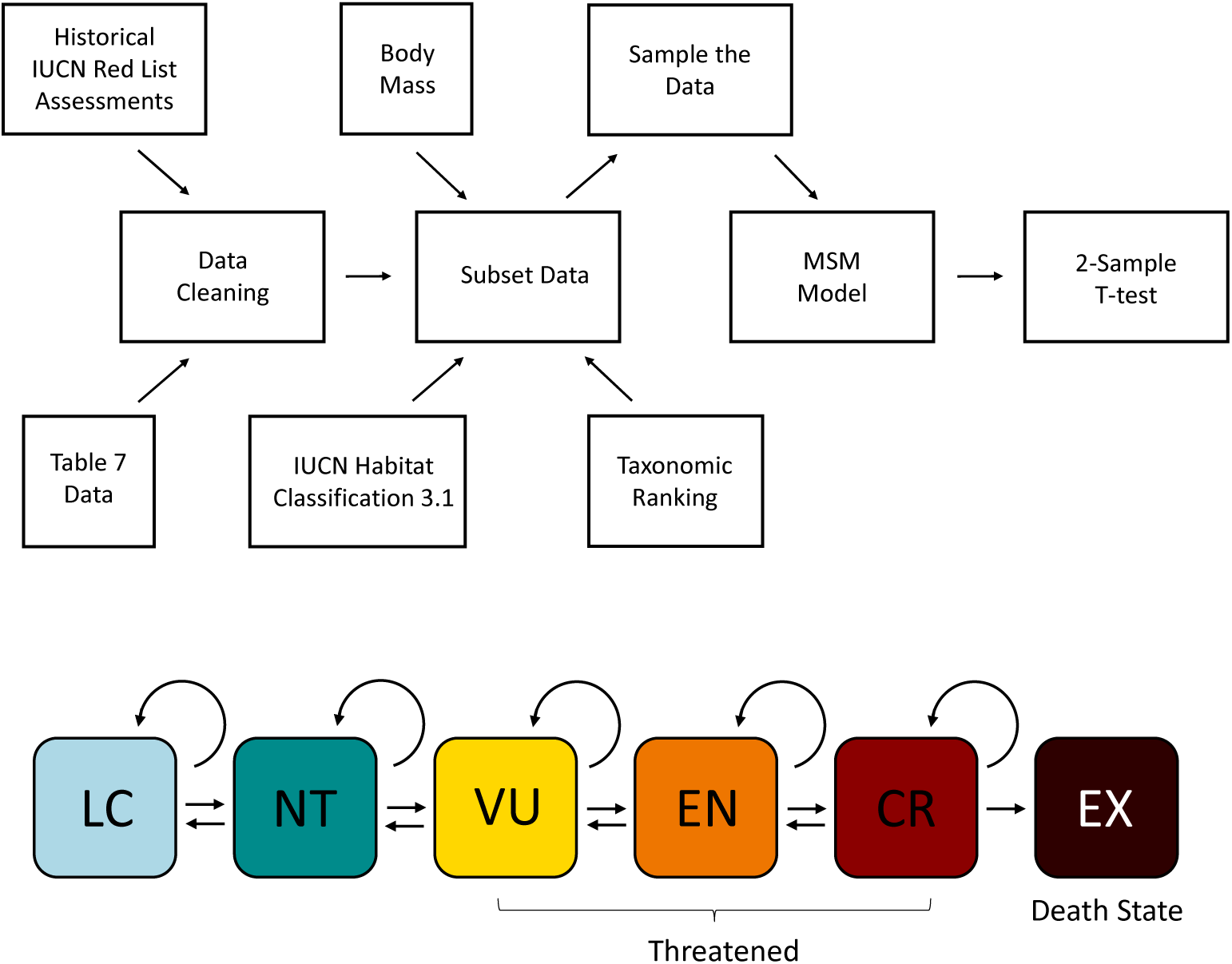
Conceptual overview of key methods components. Above: a flowchart detailing data processing and analysis steps. First historical Red List and ‘table 7’ (validity) data combine to provide genuine transitions between Red List categories and their dates. Second, the data are optionally subset by characteristics of interest such as body size and then sampled further to allow bootstrapping. Finally the multi-state Markov model (MSM) is run on each bootstrapped sample. A T-test is used when needed to detect significant differences in extinction risk between species from different subsets that are in the same Red List category. Below: a visual representation of the transitions between Red List categories possible in the multi state Markov Model that we use. Transitions are only permitted between adjacent categories in a single step, and EX is an ”absorbing” state that species cannot transition back out of.

We repeated our analyses for various different iterations of bird data. First, the Red List was subset to birds, and taken through the same steps as carried out for the data as a whole. Second, we repeated this, but with the step to incorporate validity data skipped. Finally, conducted two further bird data analysis using Red List Index data from Birdlife International (S. H. M. Butchart et al. 2007). In one case with EW merged into EX and in the other case with CR(possibly EW) and CR(possibly EX) also merged into EX. The only cleaning step required for Red List index data was category merging and removal of DD species.

In order to study the joint effects of species attributes and Red List category on extinction risk we conducted several analyses that split the data into two subsets. In each case one subset had only species with the attribute of interest whilst the other subset only had species without the attribute of interest. We then ran both subsets of the data through the MSM model 100 times using the same bootstrapping approach as previously (see Figure 1). For each comparative analysis, we performed two-sample T-tests with Bonferroni corrections on the extinction probability at 100 years, for each Red List category, across all 100 bootstrapping runs, to test if the groups’ probabilities of extinction were significantly different from one another. In some cases we also calculated a ratio of extinction risks 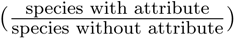 at 100 years for each threat category to be able to directly compare groups and see if an association has a positive or negative effect on extinction risk. This yielded a distribution for each threat category where values around zero indicate no effect of the attribute on extinction risk beyond those already captured by Red List category.

For the taxonomic analysis, we split the data into taxonomic groups that are well-represented in the Red List and contained enough data to model (*≥* 1500 species depending on composition). This resulted in six groups, mammals, birds, amphibians, fish, plants, and invertebrates. Reptiles and fungi are the two main groups of interest not represented due to lack of data; Reptiles included only two Extinct species and therefore the MSM parameter inference had difficulties converging, whereas only five fungi species had more than two genuine assessments with the current criteria. For each taxon group that we analysed, we split the overall data into two sections: the species belonging to that taxon, and the remaining species.

For the body mass analysis, we only looked at birds and mammals as those groups have been assessed multiple times and have comprehensive body mass data (Tobias et al. 2022; Wilman et al. 2014). We split the data in two sections based on median body mass. As there are only ten EX mammals and ten EX birds, 0.24% and 0.1% of the species respectively, the dataset was possibly too small for a reliable analysis despite the model being able to converge. To counter this, and provide a sensitivity analysis, we also modelled the probabilities of species moving not only to extinction, but to either the Critically Endangered or the Extinct category over the same time period, a weaker requirement that was met by a much larger number of species increasing the precision of the model.

For the habitat specialism analysis, we gathered habitat data from the Red List database for all viable species. Specialist species are defined as those that live in only one habitat type, whereas generalists live in more than one habitat type (as defined by Level 1 of the IUCN Habitats Classification Scheme 3.1 2020). We then modelled specialist and generalist species as separate subsets (as per Figure 1).

## 3 Results

The data handling steps produced a database of 108,007 assessments from 26,808 species, down from the original 240,816 assessments from 114,880 species. 72,930 species were lost due to only having been assessed once with the current criteria (see supplemental mats for full breakdown). Of the final data, 86% of species were animals (23,149), of which 43% (9976) were birds.

As expected for the analyses covering all species with valid transition data, probability of extinction increases both with time and with each increase in severity of Red List category (up-listing). This increase in extinction risk from up-listing is non-linear, with a super-linear growth in extinction risk through Red List categories from LC to CR. Probability of extinction at 100 years is 0.15 for CR species. The 95% confidence intervals for each Red List category do not overlap, and widen as the categories go from least to most threatened (Figure 2). Andermann *et al*. (2021) calculated a overall extinction rate across all bird species of 6.98 *×* 10*^−^*^4^ at 100 years. Using our figures, that number comes out to 2.1 *×* 10*^−^*^2^ over 100 years across all species in our data, or 2.7 *×* 10*^−^*^2^ across all non-DD extant species currently on the Red List.

**Figure 2:**
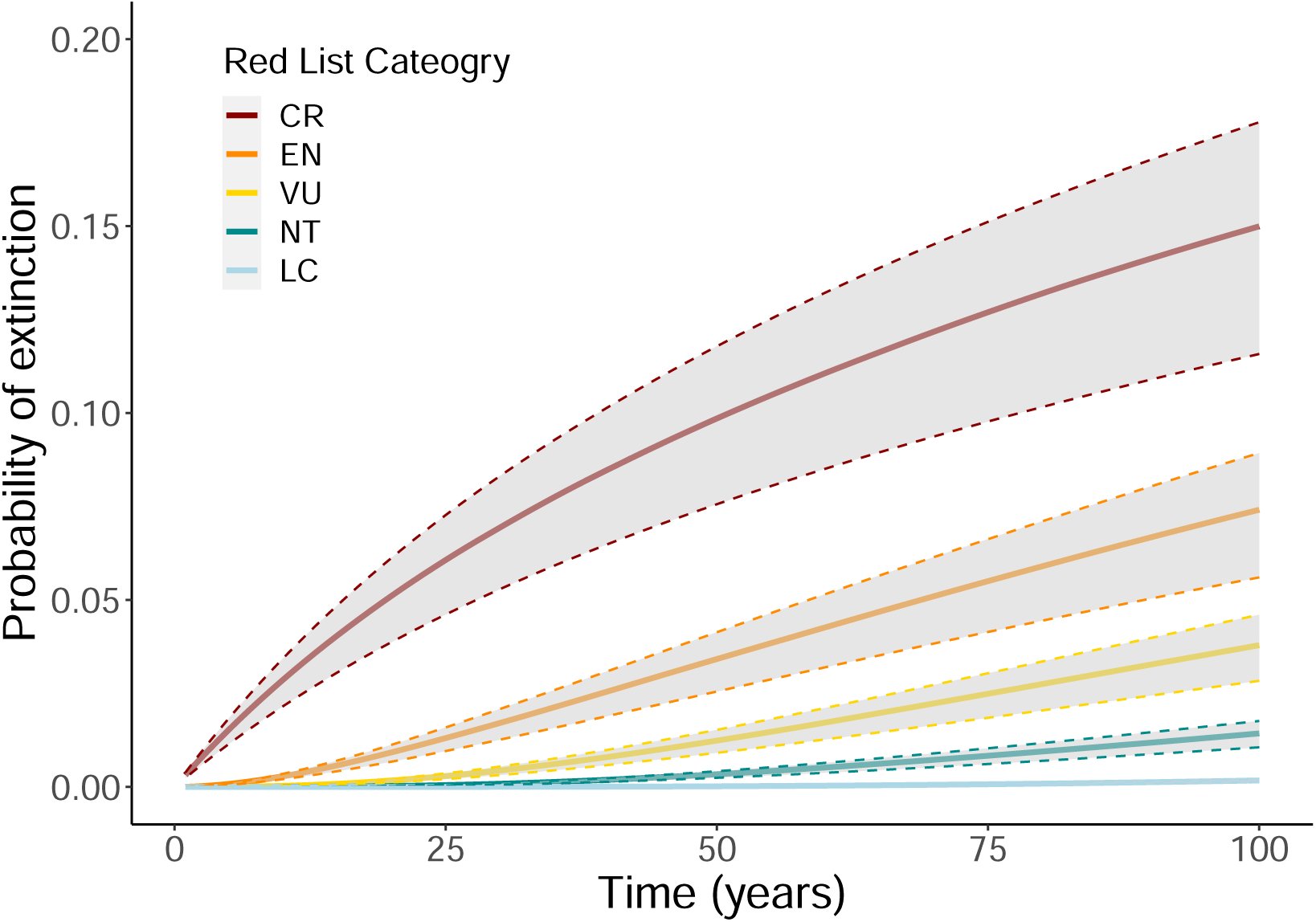
Probability of a species going Extinct over the next 100 years given its current assessed Red List category. The greyed uncertainty bounds are the 95% confidence intervals based on boot-strapping by sampling with replacement. Values are calculated by using multi-state Markov models to predict future category changes given past Red List category changes.

For the birds only analyses, extinction risk was lower than that of all species across all categories except for CR RLI species treated pessimistically (Table 1). There was very little difference in extinction risk between the Red List data with validity data incorporated and the Red List data without (Table 1). The Red List Index data produced lower extinction risk estimates than the Red List data by up to an order of magnitude, for all categories except EN and CR. (Table 1) For EN species the pessimistic Red List Index data (with CR(possibly EW) and CR(possibly EX) species treated as EX) had the same extinction risk as the Red List data. Additionally, for CR species the base Red List Index data had the same extinction risk while the pessimistic data produced an almost doubled risk of extinction (see Table 1). Extinction risk estimates were similar enough between the Red List data (both corrected and not) and the Red List Index data that there was substantial overlap across the 95% uncertainty bounds (see Supplementary Materials for significance testing).

**Table 1:**
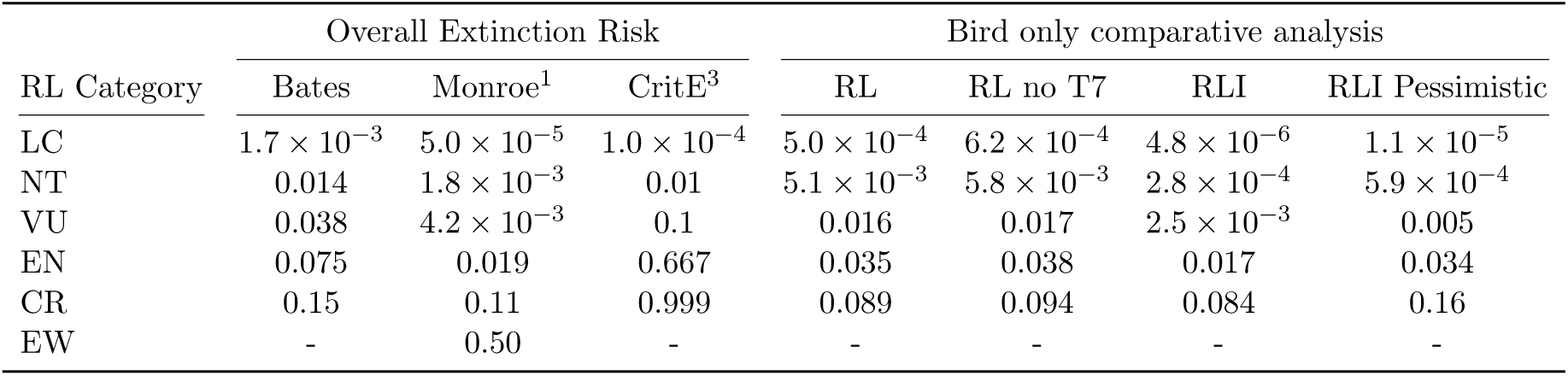
Median extinction risk for species in a given Red List assessment category at 100 years, rounded to two decimal places. ^1^Values extrapolated from the transition matrix given in Monroe et al. 2019 (note that they modelled EW seperately). ^2^ Values taken from the 100 year extrapolation of Criterion E extinction risk values in Mooers et al. 2008. For the Birds only analysis, RL indicates Red List data, while RLI indicates Red List Index data. The ”no T7” analysis is where the validity data was not used to correct the base Red List data, while the ”pessimistic” analysis reassigns CR(Possibly EX) and CR(Possibly EW) as EX.

For the analysis of body mass, splitting mammals at the median body weight of 91.3g led to a heavy group of 6116 assessments across 1917 species, and a light group of 6511 assessments across 2146 species, while splitting birds at their median body weight of 38.1g led to a heavy group with 21193 assessments from 3043 species, and a light group with 32900 assessments across 4816 species. Extinction risk increased across the categories, for both groups. For mammals, heavier species had a higher extinction risk in each category except CR, and for birds the lighter species had a higher extinction risk in all categories.

However for both groups (and mammals especially), the difference between the two body mass categories was small with a large amount of overlap (Figure 3; Table 5 Supplementary Materials). When the probability of a species becoming either CR or EX was modelled (instead of probability of extinction), mammals showed a much larger difference between the lighter vs heavier species across all categories. Birds however, showed the opposite pattern to our results for probability of extinction, with heavier birds having a higher probability of becoming CR or EX than lighter species. Although the difference is small, this pattern was true for all Red List categories (see Figure 3).

**Figure 3:**
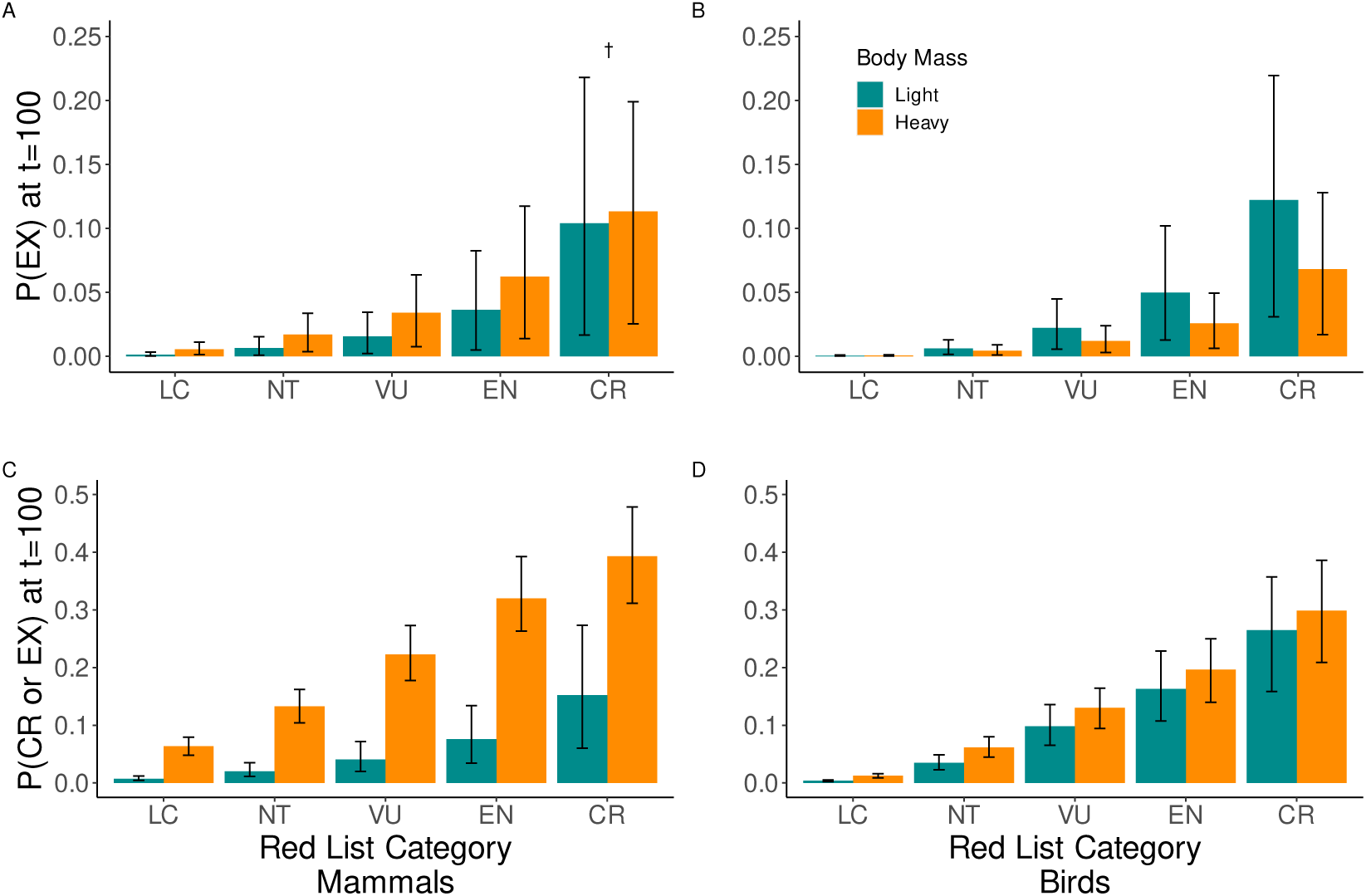
Risk of extinction (A and B) or becoming/remaining Critically Endangered or Extinct (C and D) in 100 years, for both mammals and birds. All groups were split by body mass at the median (38.1g for birds, 91.3g for mammals). Bars indicate 95% certainty intervals. The *†* indicates where the light and heavy species did not have significantly different extinction risks (p*<*0.001, Bonferroni corrected two sample T-test).

For the taxonomic group analysis, species were divided into birds (9976 species/37% of total species), mammals (4232/16%), amphibians (2738/10%), invertebrates (2222/8%), and plants (3524/13%). Plant and bird species were uniformly at lower risk of extinction compared to other species in the same Red List category. Fish and invertebrates conversely had a uniformly higher risk of extinction compared to other species in the same Red List category. Least Concern mammal species had a higher risk of extinction than other LC species, but mammals species in all other Red List categories had a lower risk of extinction than other species in the same Red List category. Amphibian species were generally at slightly lower risk of extinction compared to other species in the same Red List category, but the difference was only statistically significant for those in the three threatened categories (Figure 4).

**Figure 4:**
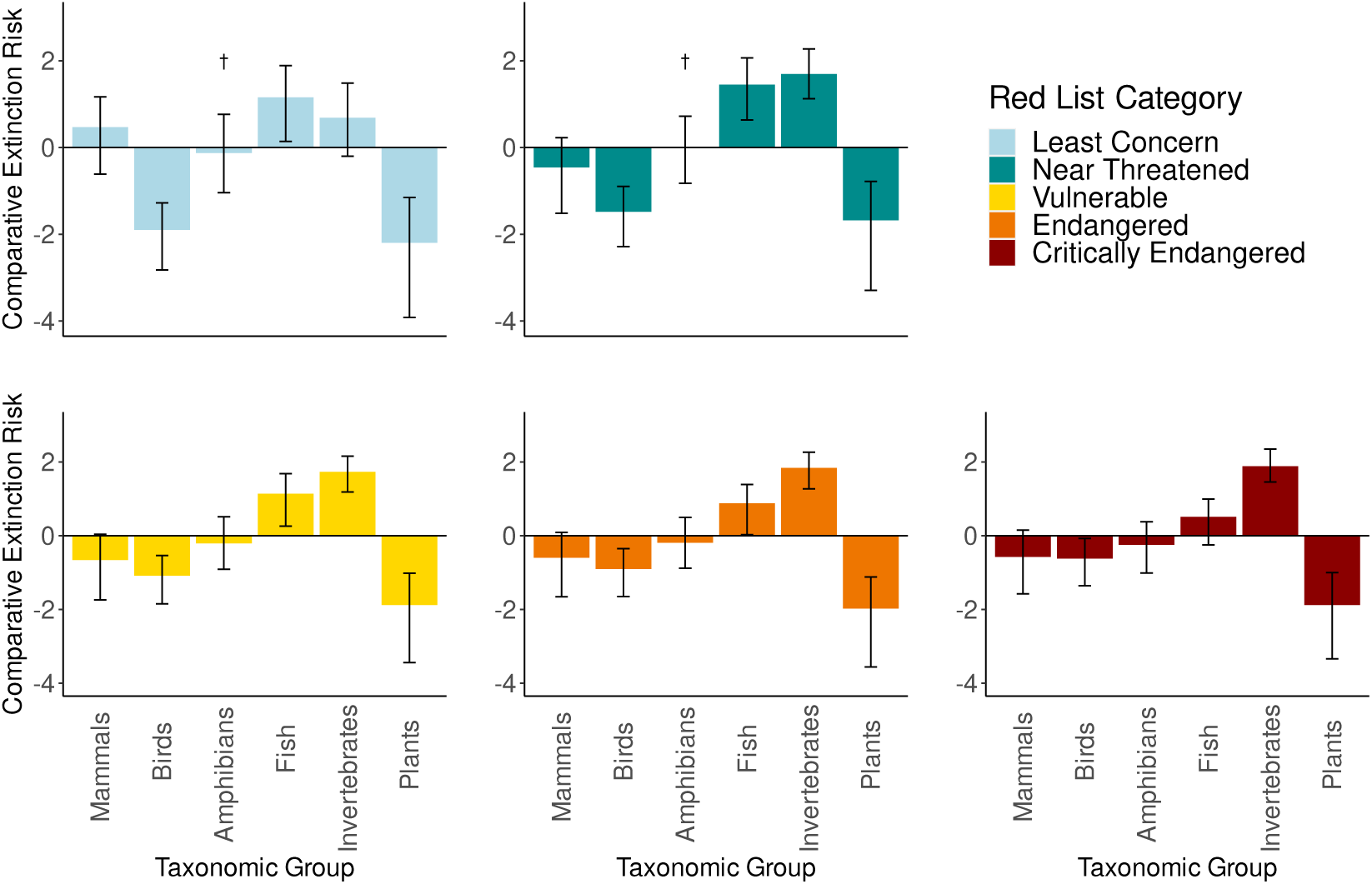
Comparative extinction probabilities across Red List categories and taxon groups at 100 years. Each bar describes the log ratio of extinction probabilities for species in the given taxa vs all other species. A negative value shows that the given taxon has a lower probability of extinction and a positive value shows that the given taxon has a higher probability of extinction. Error bars describe 95% confidence intervals. The *†* indicates where the light and heavy species did not have significantly different extinction risks (p*<*0.001 with Bonferroni correction).

Specialist species were at a higher risk of extinction than generalist species in the same Red List category. This was true for all five categories. The difference between probability of extinction for the specialist and generalist species increased with the severity of the Red List category. A habitat specialist species that is EN has a similar extinction risk to a CR habitat generalist species, and additionally a CR habitat specialist has roughly twice the risk of extinction as a CR habitat generalist (Figure 5).

**Figure 5:**
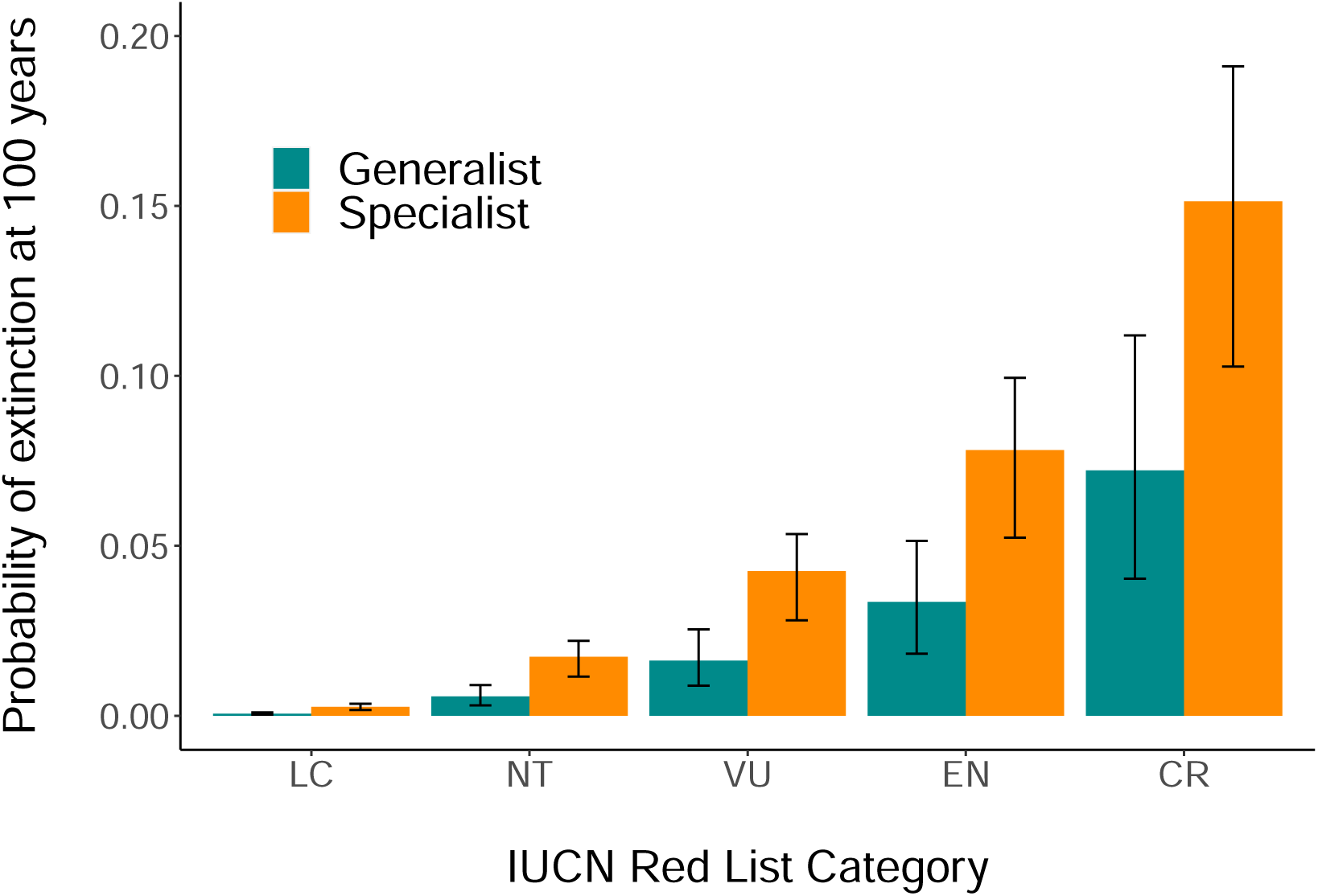
Extinction risk in 100 years across the Red List categories for both specialist and generalist species, where specialists only occupy one habitat (as per the Level 1 of the IUCN Habitat classification scheme 2020) but generalists occupy several. The extinction probabilities were significantly different at each Red List category.

## 4 Discussion

In this study we introduced a new analysis pipeline that draws from both historical IUCN Red List assessments and validity data to predict probabilities of extinction from IUCN Red List category and other factors. To the best of our knowledge, this manuscript is the first attempt to generate extinction probabilities across all assessed taxa using Markov modelling and historical IUCN assessments. Previous work in this space used data for birds only, as it is the only RLI taxon group with more than two assessments per species, and has readily available species’ generation lengths (Andermann et al. 2021; Monroe et al. 2019) . Our method enabled an investigation of all taxa present using validity data to account for non-genuine status changes, and compared this approach with using the ”gold-standard” Red List Index data. Our analyses also enabled us to test whether species attributes have effects on extinction risk beyond those already captured by the Red List category species are placed in. We found that habitat specialists are more at risk than habitat generalists in the same Red List category. Fish and invertebrates tend to be more at risk than mammals, birds, and plants in the same Red List category. The effects of body mass were mixed but generally show larger body size leads to greater risk.

Our predicted probabilities of extinction based on the whole Red List data set showed, as expected, a clear increase over time and with increasing severity of Red List categories. Our results for birds were comparable to those achieved with only Red List Index screened data, particularly for the Critically Endangered and Endangered categories. These are the most important categories to have robust estimates for as they reflect which species are most at risk and therefore used disproportionately to inform conservation decisions. This suggests that even without the additional checks and back-casting present in Red List Index data, the results gained from modelling Red List data approximate real world extinction risks reasonably well.

The results of our body mass analyses were mixed, with differences between mammals and birds. Heavy species had a higher probability of becoming either CR or EX compared to lighter species starting in the same Red List category. This result was very strong for mammals, and less strong for birds, but still with statistical significance. Heavy species also had a higher probability of extinction for mammals, but a lower probability of extinction for birds. It was not our aim here to compare extinction risk directly between mammals and birds, but rather to compare the residuals between those already in the same Red List category. The reasons for our results may therefore be a quite intricate combination of factors that differ between taxa and are not incorporated into the criteria. Small data pools inhibited our ability to do more granular analyses on patterns of risk; for example previous studies have identified a bimodal pattern of risk that would be missed by our analyses (Ripple et al. 2017). The patterns may also be driven by non-taxon specific effects driven by the different patterns of body mass across taxon groups, for example mammals have a larger median body mass than birds (91.3g vs 38.1g). We hope that future work will investigate ways to incorporate the differences we expose in future analyses of extinction risk.

Our taxonomic analyses give insights into how taxonomic differences may interact with the Red List assessment criteria. Plants from any Red List category appear to have a lower probability of extinction compared with other taxa in the same category. In contrast, fish and invertebrates have an understated risk of extinction compared to other taxa from the same Red List category. Mammals and birds, two highly charismatic and often focal groups to conservation efforts, both show lower extinction risk compared to non-mammal and non-bird species in the same Red List category. It may be that the Red List criteria threshold values do not carry the same implications for extinction risk across taxa, or that some taxa are more likely to use different assessment criteria e.g. plants tend to be assessed using Criterion B (geographic range). The differences are also likely to be due to differing conservation attention and monitoring across groups (Clark and May 2002; Donaldson et al. 2016), especially in those that are under-sampled where there is likely bias in the species contained in the data used here. It is important that we understand how the Red List may assess taxa differently, as groups that have an understated extinction risk given their Red List category are at risk of being overlooked for much needed conservation attention.

Our habitat specialism analyses show that specialist species have a higher risk of extinction than generalist species in the same Red List category. The effect is strong with Endangered habitat specialists having similar extinction risk to Critically Endangered habitat generalists. A body of previous work has shown that specialist species are more at risk of extinction in general (e.g. McKinney 2003; Fisher et al. 2003; Julliard et al. 2004; Kotze and O’Hara 2003; Keinath et al. 2017). Some of this work has been based on the IUCN Red List statuses as an indicator of extinction risk, and suggests that the current criteria do partially account for the increased risk to habitat specialist species. Our results show that habitat specialists are likely worse off than their IUCN Red List category suggests, whilst habitat generalists are better off. This means that the already reported risk of extinction for habitat specialists may be a lot worse that previously suspected.

Habitat generalist species are more able to adapt to geographic changes across space and time than their specialist counterparts (Keinath et al. 2017; McKinney and Lockwood 1999). As such, it is likely that the current geographic range of habitat generalist species is much smaller than the potential range they could occupy based on habitat suitability, as evidenced by the prevalence of habitat generalists as invasive species (Marvier et al. 2004). Conversely, the present geographic range of habitat specialists may be no smaller than their potential geographic range. Habitat specialists are therefore in a much more precarious situation, with less potential to expand or move their range, compared to the habitat generalists. Habitat specialist’s reduced ability to bounce back in future does not seem to be explicitly accounted for in the Red List assessment criteria which are focused on current geographic range (criterion B) and population size (other criteria). There may be potential to add detail to criterion B to add consideration of possible range expansion, for example by incorporating AOH (extent of suitable habitat; Brooks et al. 2019).

Our results show a different pattern to previous extinction risk estimates built using Criterion E. Mooers et al. (2008) calculate a probability of extinction of 0.999 for Critically Endangered species at 100 years. Criterion E states that to be Critically Endangered, a species should have at least a 50% chance of extinction in 10 years or three generations whichever is longer, up to a maximum of 100 years. The value of 0.999 thus corresponds to the case where three generations is less than 10 years and might fall to 0.5 if three generation lengths of the species total to 100 years or more, as could occasionally be the case. In contrast, our probability of extinction is 0.15 at 100 years for a CR species (see Table 1). For Least Concern species however Moores *et al* calculate an extinction risk of 1*×*10*^−^*^4^, whereas our model predicts an extinction risk of 1.72*×*10*^−^*^3^. We calculate an overall extinction risk of 2.1 *×* 10*^−^*^2^ over 100 years across all species in our data, or 2.7 *×* 10*^−^*^2^ across all non-DD extant species currently on the Red List.

While Criterion E may give extinction risk estimates that are higher than observed risk across the Red List as a whole, our results are likely to underestimate extinction risk in most cases. This is because the data available only included extinctions known to occur since 1996. We require data not only on extinctions but on at least one Red List assessment of the species carried out for a period in history prior to their extinction using the criteria post hoc. Furthermore, the Red List has stringent requirements to avoid species being erroneously classified as EX, and require assessors to have no reasonable doubt that the last individual of a species has died. EX classifications require exhaustive surveys over a period of time relative to the species’ life history, and as such there can be a lag between suspecting extinction and recording it. Some assessors use CR(Possibly EX) and CR(Possibly EW) to bridge this gap. When treating possibly EX and possibly EW species as EX and EW respectively, probability of extinction doubles across all categories (Table 1). Possibly Extinct is not used across all taxa or recorded in the historical data (they are recorded as CR) so we could only investigate this category in the bird RLI, but our results suggest that wider adoption of this classification would be advantageous to biodiversity research. Our probability of extinction is calculated using all valid reassessed species, and therefore incorporates an average level of conservation attention per species. As a result, our category transitions and therefore estimates of extinction risk will be an underestimate of what would be experienced in a species receiving no conservation attention.

The unusual taxonomic breadth of our study introduces unavoidable biases. While the majority of described vertebrates have been assessed at least once (81% across all groups), other taxa are increasingly undersampled (plants 15%, invertebrates 2%, and fungi & protists 0.5%) (Table 1a; IUCN 2023). Even within the well-assessed groups, bias in terms of frequency of assessments can occur. Generally those species that are more abundant, in protected areas, or occupying areas in wealthy countries, are better documented (Martin et al. 2012; Roberts et al. 2016). For example, of the 108,007 historical assessments used in this study, 67,828 were from birds (62.8%) as they are reassessed more frequently and more comprehensively than other taxa in large part due to the work of Birdlife International.

Plants and invertebrates are harder to identify to species level, often requiring specialists to do so, and aquatic life is harder to access and assess (fish have the lowest proportion of described vertebrate species assessed at 70%). This pattern of taxonomic bias is consistent with conservation research as a whole (Clark and May 2002), where resources are spent where there is interest and institutional support. More poorly assessed groups may have a bias towards threatened species (e.g. Webb and Mindel 2015), however, over time the ratio of non-threatened species vs. threatened species assessed by the IUCN keeps increasing as the Red List initiative fills knowledge gaps (Figure 1; IUCN 2023).

One way the IUCN is looking to address these biases is via the Sampled Red List index (M Baillie et al. 2008). The SRLI is an addition to the Red List Index project that records the overall change in threat via Red List categorisation (and therefore extinction risk by proxy) over time for a representative sample of species in a given taxonomic group. The Sampled Red List Index varies in scope from closely related groups of species (e.g. Dragonflies) to much broader groupings (e.g. fish and reptiles) (Böhm et al. 2013; Clausnitzer et al. 2009; Miranda et al. 2022). However as yet no species group has two published SRLI assessments (Henriques et al. 2020). One possible future application of our research will be applying our extinction probabilities directly to indices of extinction risk such as the Red List index and EDGE2 (S. H. Butchart et al. 2004; Gumbs et al. 2023). Our probabilities of extinction enable changes in threat to be more accurately weighted, especially at higher threat levels. Our work also provides a route to correct for taxonomic bias directly by permitting probability of extinction to vary based on both taxa and Red List category. This work could lead to important changes to the species we prioritise for conservation intervention.

Our study opens the possibility of a wide range of future work refining our techniques and applying them to other aspects of extinction risk research. It allows investigation of other species attributes that may contribute to extinction risk but not currently be captured by the Red List criteria e.g. terrestrial vs. non-terrestrial species, or analysis of species’ threats. Disallowing recovery (i.e. movement into less threatened categories) in the model would allow investigation into the impacts of conservation on extinction risk presuming that recovery is primarily because of conservation actions.

The Red List does not hold all information in a machine readable format, and some is not accessible via the Red List API. For example we wanted to investigate the criteria used to assess each species, but this is stored as plain text and only for the most current assessment (we would need this information for historical assessments as well). Additionally, only some of the assessed criteria are included (e.g. if a species was CR under criterion B, but EN under Criterion A, only Criterion B would be formally reported). As such, we were unable to conduct analyses investigating if the assessment criteria used impacted a species’ extinction risk. The process of conducting this research has highlighted both the huge value of the IUCN Red List as a source of extinction risk data, but also the need for improved data management and accessibility in order to get the most out of this amazing resource and best facilitate practical conservation and conservation research efforts.

We hope that this work will empower downstream analyses to use quantitative measurements of extinction risk across multiple taxa, and pave the way to new studies of the joint effects of species attributions and Red List category on the true extinction risk for species. Ultimately we hope to contribute to the much larger aim of predicting quantitative extinction risk and the factors that affect it more accurately and for a much broader range of taxa.

## Acknowledgements

We thank Nick Isaac and Georgina Mace for useful discussions about the project plan at an early stage. Through JR and RB this study is an output of the Georgina Mace Centre for the Living Planet at Imperial College London. For the purpose of open access, the authors have applied a ‘Creative Commons Attribution (CC BY) licence to any Author Accepted Manuscript version arising. This research was carried out as part of the NERC QMEE CDT.

