## Supplemental Information for "Quantifying the IUCN Red List: Using historical assessments to calculate future extinction risk"

### 1 100 year probabilities for the comparisons with Birds

#### 1.1 Extinction risks at 100 years

| Threat_level | Bottom | Mean | Top |
|---|---|---|---|
| LC | 0.00024 | 0.00062 | 0.001 |
| NT | 0.0025 | 0.0058 | 0.0097 |
| VU | 0.0077 | 0.017 | 0.028 |
| EN | 0.017 | 0.038 | 0.06 |
| CR | 0.04 | 0.094 | 0.14 |

**(a)** 100 year probability of extinction for all Birds without correcting using validity data

| Threat_level | Bottom | Mean | Top |
|---|---|---|---|
| LC | 0.00021 | 0.00051 | 0.0009 |
| NT | 0.0021 | 0.0051 | 0.0087 |
| VU | 0.0066 | 0.016 | 0.027 |
| EN | 0.015 | 0.035 | 0.058 |
| CR | 0.039 | 0.089 | 0.15 |

**(b)** 100 year probability of extinction for all Birds using the methodology for correcting using validity data

| Threat_level | Bottom | Mean | Top |
|---|---|---|---|
| LC | 1.7E-06 | 5.5E-06 | 1.1E-05 |
| NT | 9.6E-05 | 0.00031 | 0.00061 |
| VU | 0.0009 | 0.0026 | 0.005 |
| EN | 0.0062 | 0.018 | 0.032 |
| CR | 0.029 | 0.087 | 0.14 |

**(c)** 100 year probability of extinction for the Bird RLI, with CR(PEX) treated as CR.

| Threat_level | Bottom | Mean | Top |
|---|---|---|---|
| LC | 5.3E-06 | 1.1E-05 | 2E-05 |
| NT | 0.00028 | 0.00059 | 0.001 |
| VU | 0.0025 | 0.005 | 0.0086 |
| EN | 0.018 | 0.034 | 0.055 |
| CR | 0.098 | 0.16 | 0.24 |

**(d)** 100yr probability of extinction for Bird RLI with CR(PEX) treated as EX

Statistical testing was carried out via pairwise Wilcoxon rank sum tests with continuity corrections and Bonferroni adjusted P values. All P values were below 0.05 for all assessment categories, except between the two Red List data sets. Those had a P value of 1 for all categories except LC which was 0.26.

#### 1.2 Graphs for extinction risk over 100 years

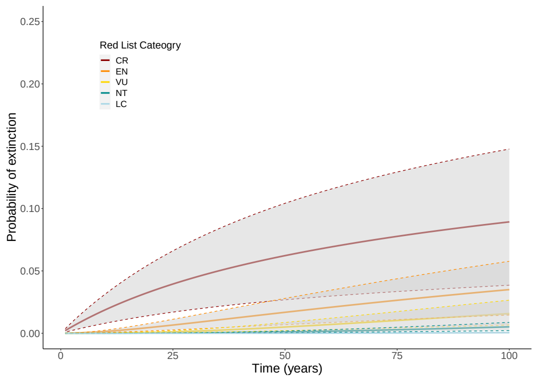


**(a)** Red List birds corrected using validity data

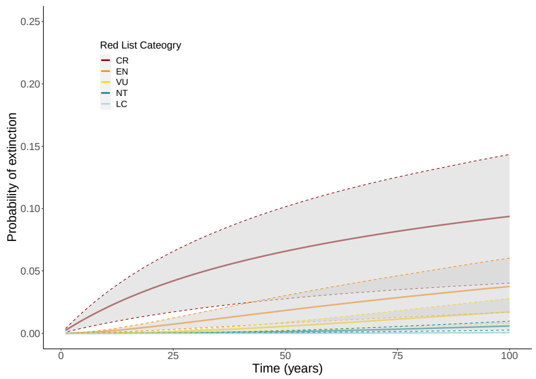


**(b)** Red List birds with no validity data corrections

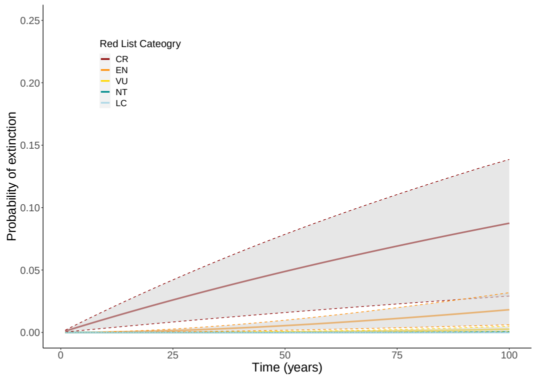


**(c)** Red List Index birds, with CR(PEX) as CR

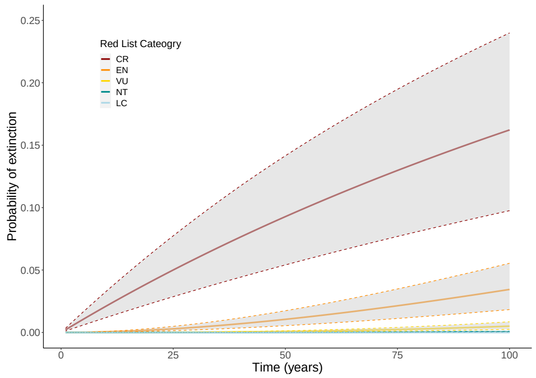


**(d)** Red List Index birds, with CR(PEX) as EX

**Figure 1:** Probability of extinction over time (with 90% confidence intervals from bootstrapping) for bird species given different input data

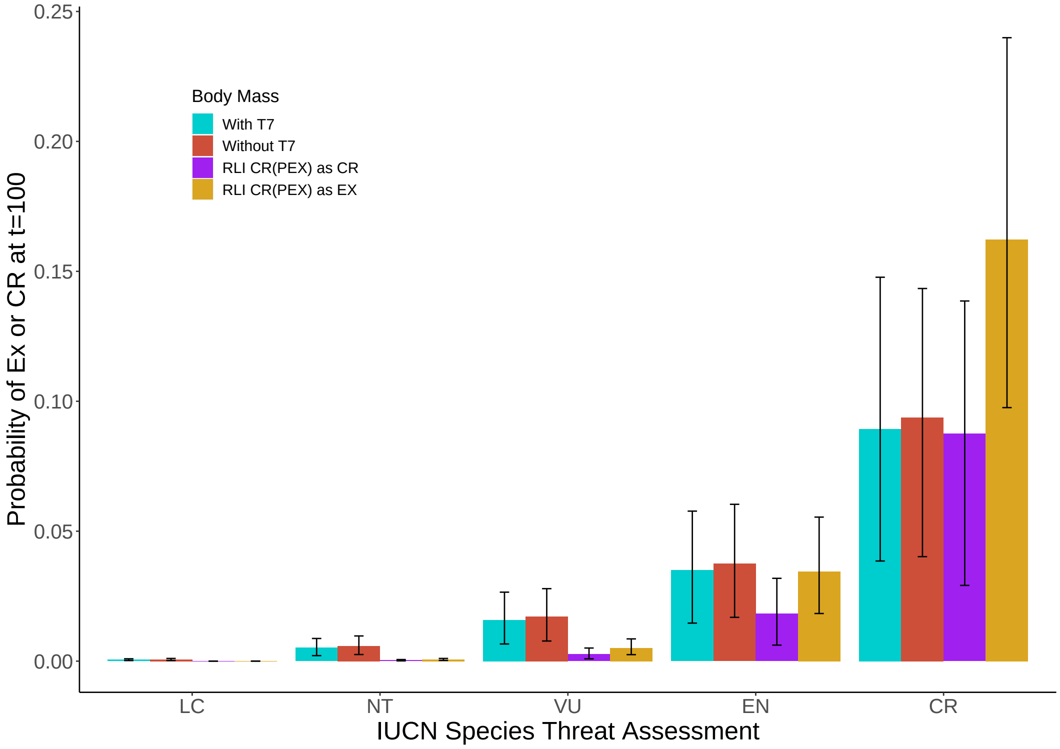


**Figure 2:** Bar chart showing probability of extinction at 100 years for various iterations of bird data. The error bars are the 95% confidence intervals from bootstrapping.

#### 2 Summary Statistics and Figures

| Assessments | Species |
|---|---|
| 1 | 72930 |
| 2 | 20104 |
| 3 | 8029 |
| 4 | 2212 |
| 5 | 933 |
| 6 | 750 |
| 7 | 787 |
| 8 | 4719 |
| 9 | 2978 |
| 10 | 874 |
| 11 | 279 |
| 12 | 128 |
| 13 | 79 |
| 14 | 44 |
| 15 | 19 |
| 16 | 13 |
| 17 | 2 |

**(a)** Number of assessments per species for the raw downloaded Red List data. Total assessments: 240816.

| Assessments | Species |
|---|---|
| 2 | 12544 |
| 3 | 4181 |
| 4 | 1211 |
| 5 | 506 |
| 6 | 690 |
| 7 | 4286 |
| 8 | 2419 |
| 9 | 596 |
| 10 | 181 |
| 11 | 86 |
| 12 | 53 |
| 13 | 30 |
| 14 | 15 |
| 15 | 8 |
| 16 | 2 |

**(b)** Number of assessments per species for the Red List data once cleaned. Total assessments: 108007

**Table 2:** Breakdown of the number of species assessed a given number of times both before and after cleaning steps

| Taxon | Assessments | Species |
|---|---|---|
| Amphibian | 6316 | 2751 |
| Bird | 67828 | 10025 |
| Fish | 5712 | 2675 |
| Invertebrate | 4967 | 2234 |
| Mammal | 13055 | 4257 |
| Plant | 7345 | 3537 |
| Reptile | 2772 | 1323 |

**Table 3:** Distribution of species/assessments per taxon group in taxonomic analysis (post-cleaning). Fungi are not present due to having too few species to model and not fitting into other taxon groups.

| | Taxon | No. species |
|---|---|---|
| Kingdom | Animalia | 73941 |
| | Chromista | 15 |
| | Fungi | 281 |
| | Plantae | 40011 |
| Animalia | Annelida | 234 |
| | Arthropoda | 12509 |
| | Chordata | 51093 |
| | Cnidaria | 884 |
| | Echinodermata | 372 |
| | Mollusca | 8831 |
| | Nemertina | 6 |
| | Onychophora | 11 |
| | Platyhelminthes | 1 |
| Cordates | Actinopterygii | 18099 |
| | Amphibia | 6649 |
| | Aves | 11123 |
| | Cephalaspidomorphi | 37 |
| | Chondrichthyes | 1106 |
| | Mammalia | 6197 |
| | Myxini | 76 |
| | Reptilia | 7796 |
| | Sarcopterygii | 10 |
| Plants | Andreaeopsida | 2 |
| | Anthocerotopsida | 2 |
| | Bryopsida | 171 |
| | Charophyaceae | 11 |
| | Chlorophyceae | 1 |
| | Cycadopsida | 324 |
| | Florideophyceae | 58 |
| | Ginkgoopsida | 1 |
| | Gnetopsida | 97 |
| | Jungermanniopsida | 85 |
| | Liliopsida | 7076 |
| | Lycopodiopsida | 90 |
| | Magnoliopsida | 30680 |
| | Marchantiopsida | 13 |
| | Pinopsida | 829 |
| | Polypodiopsida | 562 |
| | Polytrichopsida | 1 |
| | Sphagnopsida | 6 |
| | Takakiopsida | 1 |
| | Ulvophyceae | 1 |

**(a)** Number of species in various taxon groups for the raw downloaded Red List data.

| | Taxon | No. species |
|---|---|---|
| Kingdom | Animalia | 23149 |
| | Fungi | 6 |
| | Plantae | 3524 |
| Animalia | Annelida | 2 |
| | Arthropoda | 1407 |
| | Chordata | 20927 |
| | Cnidaria | 228 |
| | Mollusca | 583 |
| | Nemertina | 1 |
| | Onychophora | 1 |
| Cordates | Actinopterygii | 2272 |
| | Amphibia | 2738 |
| | Aves | 9976 |
| | Cephalaspidomorphi | 10 |
| | Chondrichthyes | 381 |
| | Mammalia | 4232 |
| | Myxini | 1 |
| | Reptilia | 1315 |
| | Sarcopterygii | 2 |
| Plants | Anthocerotopsida | 1 |
| | Bryopsida | 5 |
| | Cycadopsida | 185 |
| | Gnetopsida | 1 |
| | Jungermanniopsida | 1 |
| | Liliopsida | 464 |
| | Lycopodiopsida | 7 |
| | Magnoliopsida | 2389 |
| | Pinopsida | 451 |
| | Polypodiopsida | 19 |
| | Sphagnopsida | 1 |

**(b)** Number of species in various taxon groups for the cleaned Red List data.

**Table 4:** Number of species in various taxon groups both before and after cleaning steps. Number of species dropped from 114880 to 26808 meaning a loss of 88072 species.

#### 3 T-test stats

All p-values are corrected with the Bonferroni correction for that group

**Specialist vs Generalist:**
LC: p-value = 6.561768e-78 t = -39.926, df = 140.64 sample estimates: mean of x mean of y 0.0006065607 0.0024710715
NT: p-value = 1.348394e-75 t = -32.709, df = 174.27 sample estimates: mean of x mean of y 0.005623387 0.016268044
VU: p-value = 6.115809e-72 t = -29.777, df = 185.76 sample estimates: mean of x mean of y 0.01576356 0.04010281
EN: p-value = 5.584716e-68 t = -27.464, df = 193.5 sample estimates: mean of x mean of y 0.03216949 0.07434535
CR: p-value = 1.990344e-61 t = -24.608, df = 197.91 sample estimates: mean of x mean of y 0.06977407 0.14327181

**Body Mass:**

| Category | Birds | Birds CR/EX | Mammals | Mammals CR/EX |
|---|---|---|---|---|
| LC | 3.208066e-04 | 3.491935e-77 | 6.008569e-32 | 5.268982e-90 |
| NT | 6.012058e-05 | 5.413992e-54 | 5.412379e-23 | 1.395938e-104 |
| VU | 3.663003e-14 | 3.088851e-23 | 4.027876e-18 | 3.880620e-113 |
| EN | 1.188080e-15 | 2.996111e-11 | 3.602679e-10 | 4.307176e-97 |
| CR | 1.966997e-15 | 4.818723e-05 | 1.000000e+00 | 1.769123e-53 |

**Table 5:** p-values adjusted with the Bonferroni correction, comparing probability of extinction at 100 years between heavy and light species for a given category/origin dataset.

**Birds**
LC: t = 4.4365, df = 176.82 sample estimates: mean of x mean of y 0.0005685566 0.0004130519
NT: t = -4.8191, df = 184.07 sample estimates: mean of x mean of y 0.004311120 0.006029763
VU: t = -8.8388, df = 157.35 sample estimates: mean of x mean of y 0.01185781 0.02218498
EN: t = -9.423, df = 155.17 sample estimates: mean of x mean of y 0.02575060 0.04971442
CR: t = -9.2109, df = 175.26 sample estimates: mean of x mean of y 0.06817236 0.12215168

**Birds CR/EX**
LC: t = 41.291, df = 135.09 sample estimates: mean of x mean of y 0.012404128 0.003855245
NT: t = 22.857, df = 181.38, sample estimates: mean of x mean of y 0.06160588 0.03492089
VU: t = 11.753, df = 198 sample estimates: mean of x mean of y 0.13020063 0.09811394
EN: t = 7.5589, df = 197.19 sample estimates: mean of x mean of y 0.1966350 0.1631776
CR: t = 4.8607, df = 193.94 sample estimates: mean of x mean of y 0.2986046 0.2648532

**Mammals**
LC: t = 16.359, df = 128.91, sample estimates: mean of x mean of y 0.005515834 0.001256227
NT: t = 12.1, df = 158.71, sample estimates: mean of x mean of y 0.017006260 0.006392916
VU: t = 10.269, df = 164.32 sample estimates: mean of x mean of y 0.03407228 0.01557818
EN: t = 7.2338, df = 161.08, sample estimates: mean of x mean of y 0.06219133 0.03627353
CR: t = 1.137, df = 126.49 sample estimates: mean of x mean of y 0.1131935 0.1040481

**Mammals CR/EX**
LC: t = 63.876, df = 114.37 sample estimates: mean of x mean of y 0.063750661 0.007091297
NT: t = 64.247, df = 140.2 sample estimates: mean of x mean of y 0.13295623 0.01997112
VU: t = 62.658, df = 159.89 sample estimates: mean of x mean of y 0.22268124 0.04062623
EN: t = 51.209, df = 153.48 sample estimates: mean of x mean of y 0.31998626 0.07604886
CR: t = 28.974, df = 115.38 sample estimates: mean of x mean of y 0.3930534 0.1523343

### 4 Supplemental Methodology

Table 7 is an annual report of whether the reported transitions for that year were for genuine or non-genuine reasons. A transition for genuine reasons reflects a genuine change in the threat status of that species. a non-genuine transition is where only the knowledge of the level of threat has changed, not the level itself (e.g. taxonomic changes). Table 7 is provided as PDF documents and is available from 2007 onwards. It does not contain the reason for a transition being deemed genuine or non-genuine, nor does it contain information on how many prior assessments may be affected in the case of a non-genuine transition.

For our analyses, we used the transitions available from Table 7 (hereafter 'validity data') to label the first assessment of a pair as valid or invalid. For example, a species transitioning from EN to CR for non-genuine reasons would have the EN assessment labelled as invalid. Not all transitions are included in the validity data, and it does not include assessments from before 2007 or the most current assessment of a species. From our labelled data, the proportion of the assessments from each Red List category labelled as invalid was calculated. The proportions of each category with invalid assessments was then used to randomly assign either a valid or invalid label to those assessments without labels. This allowed us to use the validity data as a proxy for backcasting across all species rather than just a smaller subset.

Data deficient species are species for whom not enough data was available to have a category assigned to them. As such they are automatically invalid for use in our analysis and were marked as such. Additionally, any species marked as EX where there was later an extant assessment was labelled invalid, as due to the strict requirements for a species to be assessed as EX it must have been a false EX assignment.

### 5 Extinct in the Wild testing

| Threat_level | Bottom | Median | Top |
|---|---|---|---|
| LC | 0.00158 | 0.00208 | 0.00251 |
| NT | 0.0132 | 0.0168 | 0.0200 |
| VU | 0.0357 | 0.0445 | 0.0515 |
| EN | 0.0701 | 0.0865 | 0.101 |
| CR | 0.137 | 0.171 | 0.195 |

**(a)** 100 year probability of extinction for all Red List species when EW is treated as EX (rounded to 3sf)

| Threat_level | Bottom | Median | Top |
|---|---|---|---|
| LC | 0.00176 | 0.00223 | 0.00269 |
| NT | 0.0138 | 0.0171 | 0.0207 |
| VU | 0.0354 | 0.0445 | 0.0533 |
| EN | 0.0652 | 0.0827 | 0.0997 |
| CR | 0.127 | 0.163 | 0.193 |

**(b)** 100 year probability of extinction for all Red List species when EW is treated as CR (rounded to 3sf)
